# Vaccine-induced antigenic drift of a human-origin H3N2 Influenza A virus in swine alters glycan binding and sialic acid avidity

**DOI:** 10.64898/2025.12.10.693614

**Authors:** Matias Cardenas, Pradeep Chopra, Brianna Cowan, C. Joaquin Caceres, Tavis K. Anderson, Amy L. Baker, Daniel R. Perez, Geert-Jan Boons, Daniela S. Rajao

**Author notes:** Department of Veterinary Microbiology and Preventive Medicine, College of Veterinary Medicine, Iowa State University, Ames, IA, USA.

## Abstract

Interspecies transmission of human influenza A viruses (FLUAV) to swine occurs frequently, yet the molecular factors driving adaptation remain poorly understood. Here we investigated how vaccine-induced immunity shapes the evolution of a human-origin H3N2 virus in pigs using an *in vivo* sustained transmission model. Pigs (seeders) were vaccinated with a commercial inactivated swine vaccine and then infected with an antigenically distinct FLUAV containing human-origin HA/NA. Contact pigs were introduced two days later. After 3 days, seeder pigs were removed, and new contacts introduced. This was repeated for a total of 4 contacts. Sequencing of nasal swab samples showed the emergence of mutations clustered near the HA receptor binding site, enabling immune escape and abolishing binding to N-glycolylneuraminic acid. Mutant viruses recognized α2,6-sialosides with 3 N-acetyllactosamine repeats, which are rare in swine lungs, while the parental virus bound structures with a minimum of 2 repeats. Adaptative HA mutations enhanced avidity for α2,6-linked sialic acid, likely compensating for the low abundance of extended glycans. Notably, residues outside the canonical HA binding pocket contribute to glycan binding, suggesting a trade-off between receptor breadth and avidity. These findings show that non-neutralizing immunity promotes viral adaptation by fine-tuning receptor engagement and immune evasion.

**SIGNIFICANCE:** Understanding how vaccination shapes influenza A virus (FLUAV) evolution across species barriers is critical for predicting and preventing zoonotic and reverse-zoonotic events. Our study demonstrates that vaccine-induced immune pressure can drive antigenic drift in a human-origin H3N2 virus, altering HA receptor binding properties that could inadvertently facilitate adaptation to swine. These changes shifted glycan specificity toward extended poly-LacNAc structures and enhanced α2,6-linked sialoglycans binding while abolishing Neu5Gc recognition. By revealing how non-neutralizing immunity fine-tunes HA–glycan interactions by engaging antigenically relevant residues in glycan binding, this work highlights vaccination as an underappreciated driver of host adaptation and viral evolution.

## INTRODUCTION

Influenza A viruses (FLUAV) circulate endemically in both humans and swine. Frequent human-to-swine transmission contributes to the genetic diversity of FLUAV in pigs [1–3]. However, the molecular determinants that lead to sustained circulation following a reverse zoonotic event remain poorly understood [4]. Reassortment and acquisition of swine-adapted internal gene segments seems to be a critical initial step in viral establishment [4–7], while subsequent adaptative mutations in or near the hemagglutinin (HA) receptor binding site (RBS) have been linked with efficient pig-to-pig transmission and long-term establishment of these novel lineages [8, 9].

Vaccination is the most effective strategy to prevent FLUAV infections in swine [10]. Inactivated commercial vaccines, widely used in swine farms, elicit strong HA-specific IgG responses [11] and lower neuraminidase (NA)-specific response [12]. Nevertheless, protection against infection is limited unless the vaccine closely matches the circulating strain [13], albeit mismatched vaccination may still decrease onward transmission and disease severity. Antibody-mediated immunity creates a selective pressure that may drive the emergence of genetic variants and that subsequently affect antigenic phenotypes [14]. Vaccine-driven antigenic evolution has been shown following immunization of pigs, with mutations arising in the HA RBS and the NA head domain [15, 16].

The HA protein binds to sialic acid (SA)-containing glycans (sialoglycans) on the host cells and therefore is one of the main host range determinants of FLUAV [17]. Avian viruses mostly bind *N*-glycans with α2,3-linked sialic acid (α2,3-SA) [18], while mammalian viruses exhibit a preference for α2,6-sialosides (α2,6-SA) [19]. Despite the presence of α2,6-SA in both humans and swine respiratory tracts [20, 21], human FLUAV still require modifications to the HA to establish sustained infection in pigs. Enhanced affinity for *N*-glycolylneuraminic acid (Neu5Gc), which is expressed in pigs [22] but not in humans [23], has been proposed as a swine-adapting modification.

However, some studies have reported no difference in Neu5Gc affinity between human and swine FLUAV [24–27]. Therefore, the impact of HA mutations following human-to-swine spillovers remains unknown. Moreover, it is unclear whether vaccine-driven changes in the HA RBS affect receptor affinity and/or could facilitate adaptation.

Here we examined how vaccine-driven evolution shapes human H3N2 adaptation to swine. For this purpose, vaccinated pigs were inoculated with a human-origin reassortant virus that was previously shown to be transmissible in pigs after acquiring a mutation near the HA RBS (A138S) [28, 29]. Serial transmission among vaccinated pigs led to loss of the A138S mutation and acquisition of new HA mutations surrounding the RBS (V186G and F193Y), while no changes were observed in non-vaccinated animals. The vaccine-induced HA changes (V186G and F193Y) reduced antibody recognition and reshaped glycan interactions. The V186G and F193Y mutations abolished binding to Neu5Gc and promoted binding to structures with at least 3 LacNAc repeats while enhancing binding to α2,6-SA. This suggests that adaptation imposed the requirement for poly-LacNAc repeats, reflecting a trade-off between receptor breadth and avidity. Importantly, these results suggest that residues outside the canonical sialic acid-binding pocket likely contribute to multivalent interactions within the HA trimer. Our results provide evidence that non-neutralizing immunity can shape cross-species adaptation of human FLUAV to swine by fine-tuning glycan recognition most likely as a consequence of immune escape. This offers insights into the molecular mechanisms favoring interspecies transmission and host adaptation.

## RESULTS

### Vaccination reduced but did not inhibit hVIC/11-A138S replication *in vivo*

Vaccination imposes strong immune pressure that drives antigenic drift of FLUAV in humans [30]. In pigs, immunization creates a similar landscape, yet its effect on viral adaptation after a spillover is poorly understood. To investigate this, we mimicked a spillover by infecting vaccinated and non-vaccinated pigs with a human-like H3N2 virus (hVIC/11-A138S) followed by four rounds of transmission **(Fig 1A)**. This virus carried 5 swine-adapted gene segments from sOH/04 coupled with the pandemic M segment from A/California/04/2009 (TRIG backbone), and the HA and NA segments from the human seasonal A/Victoria/361/2011 [29]. It also contained the A138S HA mutation (H3 numbering) that enables pig-to-pig transmission [28]. The virus gene constellation mimicked an initial reassortment event, a critical step in early adaptation [5], allowing us to focus on HA and NA evolution. Vaccinated pigs exhibited reduced viral titers **(Fig 1B)**, with an average of 1x10^4^ TCID_50_eq/mL, compared to non-vaccinated controls **(Fig 1C)**, yet transmission still occurred. Non-vaccinated pigs supported robust viral replication, with an average of 1x10^5^ TCID_50_eq/mL across contacts. While differences in the vaccinated group were not statistically significant, a trend toward higher titers was observed in later contacts (contacts 4, C4 pigs). This effect was more apparent in bronchoalveolar lavage fluid (BALF), where hVIC/11-A138S was initially undetectable, but later appeared in the lungs of C4 pigs **(Fig 1D)**. Antibody titers remained stable throughout the study, with no significant variation between the 14 days post-boost and the time of exposure **(Fig 1E)**. Together, these results show that vaccination reduced but did not prevent infection, and increasing titers in later contacts suggest adaptation during pig-to-pig transmission.

**Fig 1.**
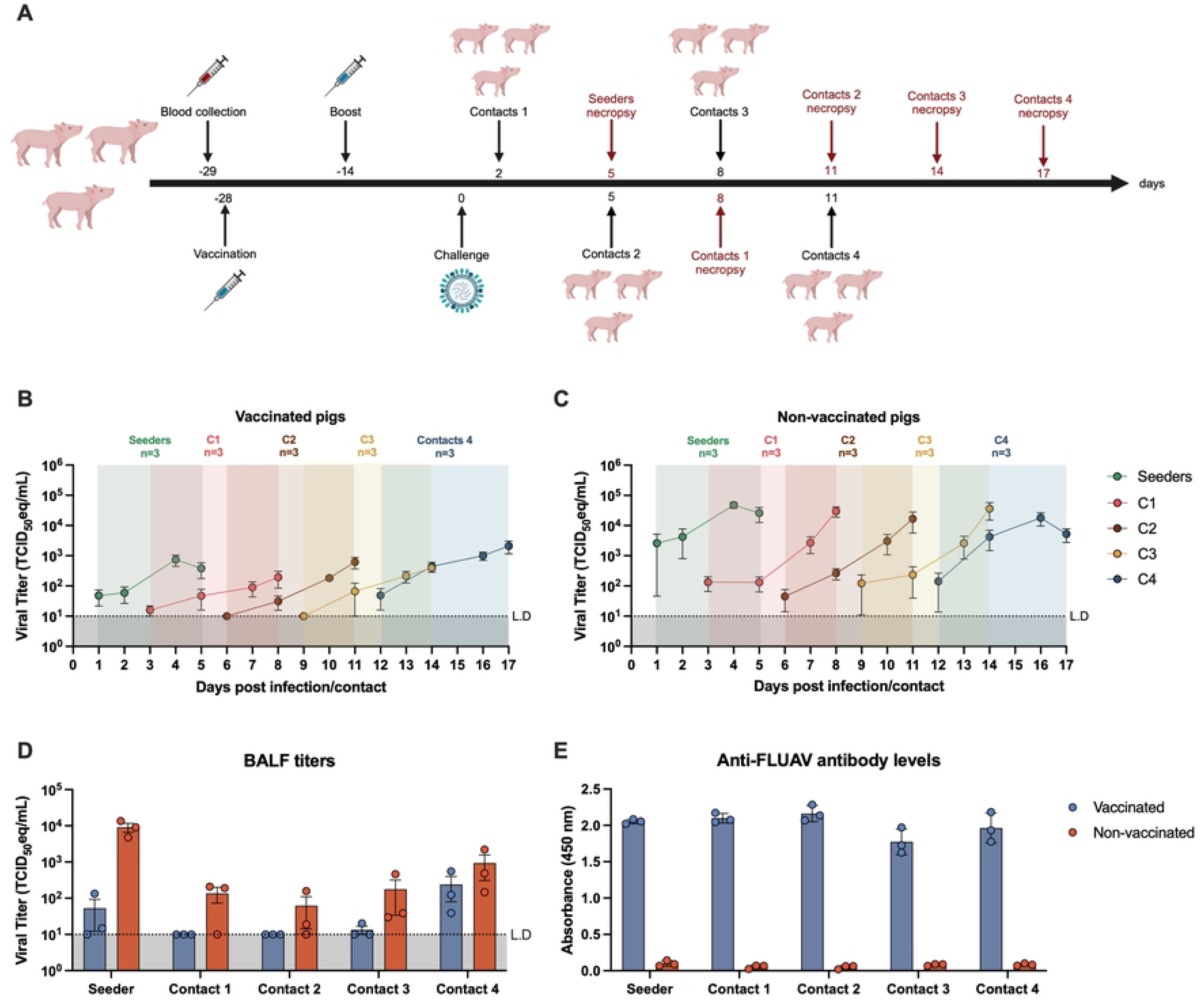
Immunization does not prevent hVIC/11-A138S infection and transmission. **a.** Animals were immunized in a prime-boost regimen. 14 days post-boost, pigs were challenged with hVIC/11-A138S (seeders). At 5 dpi, seeders pigs were euthanized, and three new pigs within the same vaccine status (vaccinated or not) were introduced as contacts (C1). This process was repeated for a total of four rounds of transmission(C1-C4). Nasal swabs were collected from vaccinated (**b**) or non-vaccinated (**c**) during the study and further titrated by RT-qPCR. Similarly, virus present in bronchoalveolar lavage fluid (BALF) was titrated by RT-qPCR (d). Anti-FLUAV antibody levels were measured for all vaccinated and non-vaccinated pigs at 14 days post-boost and the day they were placed as contacts by ELISA. Samples are presented as the mean of three pigs ± SD.

### Transmission in vaccinated animals selected for mutation in the HA gene

Nasal swab samples from vaccinated and non-vaccinated pigs with a Ct value <30 were sequenced by next generation sequencing. In vaccinated seeders, the HA A138S mutation was rapidly lost **(Fig 2A)**. Two additional RBS mutations, V186G and F193Y, emerged in C1 pigs and were transmitted to C2 contacts. V186G was transient and was quickly replaced by F193Y, which became fixed in C3 and C4 pigs **(Fig 2A)**. Structural mapping using the hVIC/11 HA crystal structure (PDB: 4O5N [31]) showed that theses residues are adjacent to the RBS but do not directly interact with sialic acid **(Fig 2B**), which interacts only with R222, N225, Y98, S137, T135, and D190. No HA changes were detected in non-vaccinated animals **(Fig 2A)**, where S138 was maintained, consistent with our previous work [32]. These findings suggest that F193Y fixation most likely resulted from vaccine-induced immune pressure.

**Fig 2.**
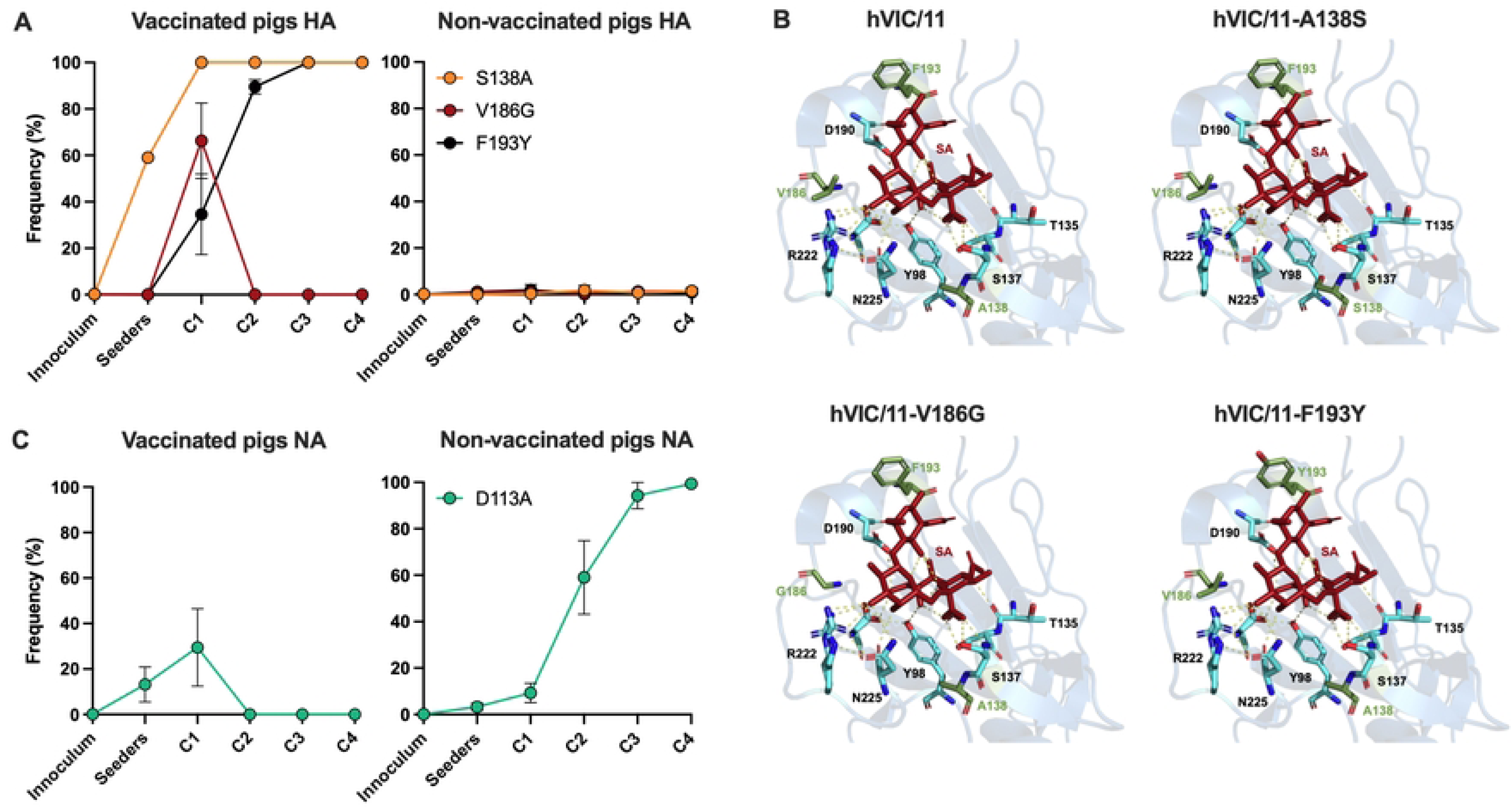
Transmission in vaccinaded pigs selected for mutations near the HA RBS. Nasal swabs collected at 1, 2, and 5 days post-inoculation (dpi) and 1, 3, and 6 days post-contact (dpc) from vaccinated and non-vaccinated pigs with a Ct <30 were sequenced and viral variants across the HA (**A**) and NA segments were identified. Major variants were defined using a threshold frequency of 0.5 (50%), while fixation was marked at a frequency of 100%. Data represent viral diversity observed in three pigs per contact group. (**C**)Cartoon representation of the hVIC/11 HA RBS showing residues 138, 186, and 193 in green and amino acids directly interacting with sialic acid in light blue. Model made using PyMOL.

In vaccinated animals, the NA D113A mutation emerged in seeder pigs but was no longer detected in C2 pigs **(Fig 2C)**. This mutation is rare among both human and swine H3N2 viruses and we previously proposed it restores the HA/NA balance disrupted by the A138S substitution in the HA [32]. The NA D113A mutation was the only change observed in non-vaccinated animals, with increased frequency over time **(Fig 2C)**. Overall, sequencing data suggest that vaccination imposed a strong immune pressure on the HA, while the NA gene remained largely conserved.

To determine the functional impact of these HA changes, we generated recombinant viruses carrying single (A138S, V186G, or F193Y), double (A138S+V186G; A138S+F193Y; or V186G+F193Y), or triple (A138S+V186G+F193Y) combinations of these mutations in the wild type HA segment from A/Victoria/361/2011 (hVIC/11), paired with or without the NA D113A mutation. Noteworthy, all these viruses contained a TRIG backbone as described above. Replication in MDCK cells was comparable among all variants and the controls (hVIC/11 and the swine-adapted sOH/04) **(S1 Fig)**, and the D113A mutation alone had no effect. Similarly, replication in differentiated human airway epithelial cells were unchanged for hVIC/11 variants except hVIC/11-F193Y, which showed reduced titers when paired with D113A **(S2 Fig)**. This indicates that vaccine-driven HA and NA changes might not affect replication capacity in humans.

### V186G and F193Y reduce antibody-mediated virus neutralization

To determine whether these mutations affected antigenicity, we tested sera from non-infected vaccinated pigs against the mutant viruses. Vaccination with a commercial swine vaccine elicited low neutralizing antibody titers against hVIC/11 (ISD50: <20, **Fig 3A)**, but higher titers against hVIC/11-A138S (ISD50: 405.4, **Fig 3B**). Neutralization titers were drastically reduced for V186G (ISD50: 131.4, **Fig 3C**) and F193Y (ISD50 49.9, **Fig 3D**) compared to hVIC/11-A138S, and this reduction extended to all double mutants (**Fig 3E-G**; A138S+V186G: 78.9 ISD50, A138S+F193Y: 73.2 ISD50, and V186G+F193Y: 86.1 ISD50). The triple HA mutant (A138S+V186G+F193Y) showed a partial increase (ISD50=239.6, **Fig 3H**) but remained below hVIC/11-A138S. The contemporary vaccine-related control A/swine/NY/01104005/2011 H3N2 (NY/11) exhibited one of the highest neutralization profiles (ISD50 650, **Fig 3I**), whereas sOH/04, an older isolate from the same lineage as NY/11, showed low reactivity (ID50 24.5, **Fig 3J**). The effect of NA D113A was context-dependent, enhancing neutralization of hVIC11 but reducing that of hVIC/11-A138S virus. Notably, when paired with HA V186G or F193Y, D113A markedly increased antibody recognition.

**Fig 3.**
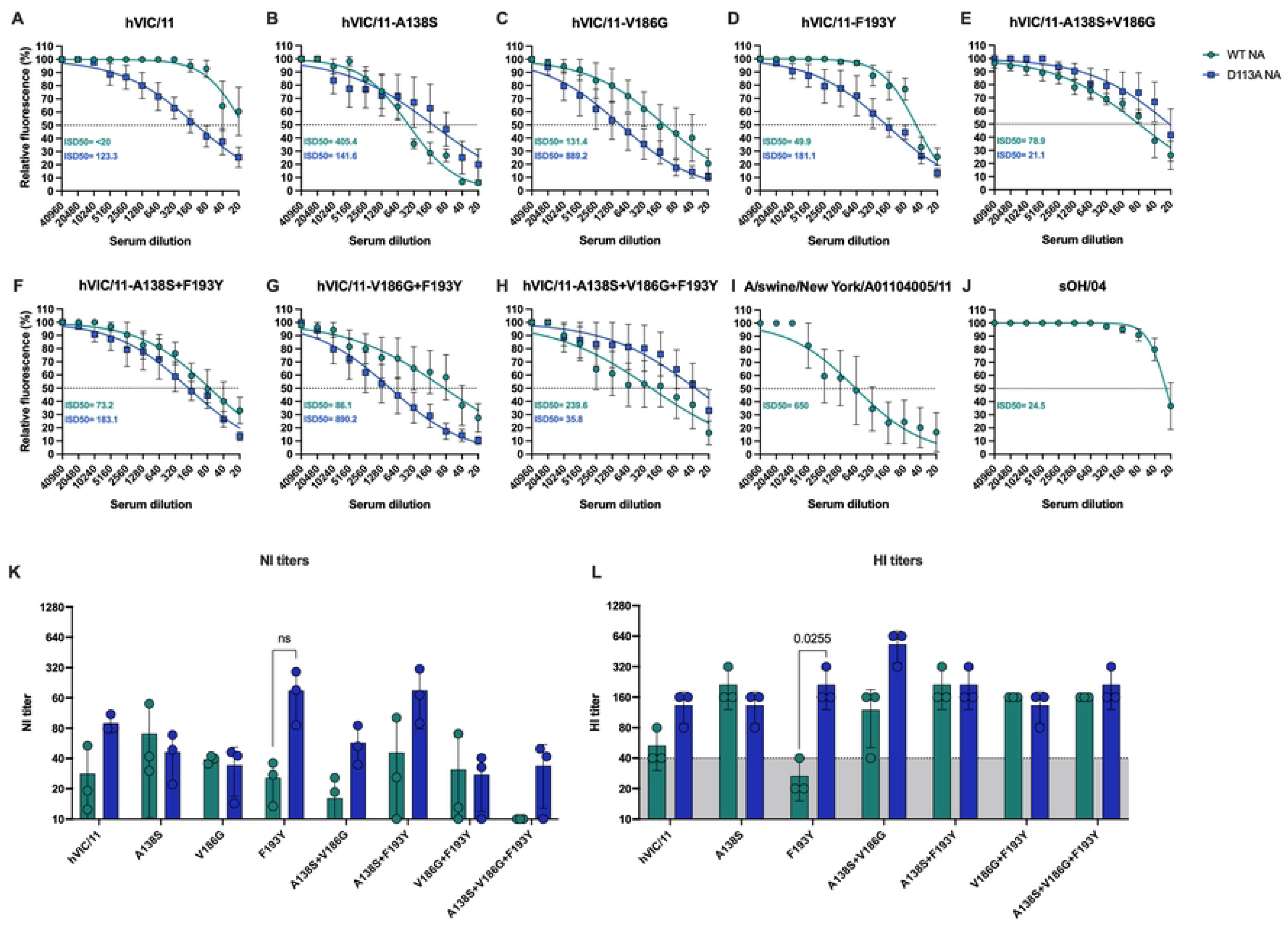
V186G and F193Y strongly reduced HA recognition by sera antibodies. Serum samples collected 14 days post boost were analyzed for the antibody response. Microneutralization assays against all hVIC/11 HA and NA variants (**a-h**), A/swine/New York/A0110405/11 (**i**) representing an antigenically similar vaccine control, and sOH/04 (**j**) were performed by incubating 100 TCID_50_ of each virus with different sera dilutions. Inhibitory titers are shown as the inhibitory sera dilution 50 (ISD50). Dotted line represents the 50% inhibition. **k.** The presence of anti-NA antibodies was assessed by evaluating NA activity using MUNANA at different sera dilutions. Neuraminidase inhibition titers (NI) are shown as the sera dilution in which 50% of NA activity was lost compared to the no serum control. l. Hemagglutination inhibition (HI) titers were determined except for hVIC/11-V186G. Dotted line represents an HI titer of 40. Data is shown as the mean of three independent experiments ± SD.

Neuraminidase inhibition (NI) assays **(Fig 3K, S3 Fig)** showed minimal effect of D113A on NI titers, except in hVIC/11-F193Y/D113A, which showed a non-significant increase compared to hVIC/11-F193Y. Hemagglutination inhibition (HI) assays **(Fig 4L)** revealed no differences between variants with or without the NA D113A mutation, except for hVIC/11-F193Y/D113A which exhibited higher titers compared to hVIC/11-F193Y (HI<40). hVIC/11-F193Y had the lowest HI titers among all mutant HAs. hVIC/11-V186G did not agglutinate chicken nor turkey red blood cells (RBCs). Overall, F193Y seems to confer an advantage under immune pressure by decreasing HA antigenicity and neutralization.

**Fig 4.**
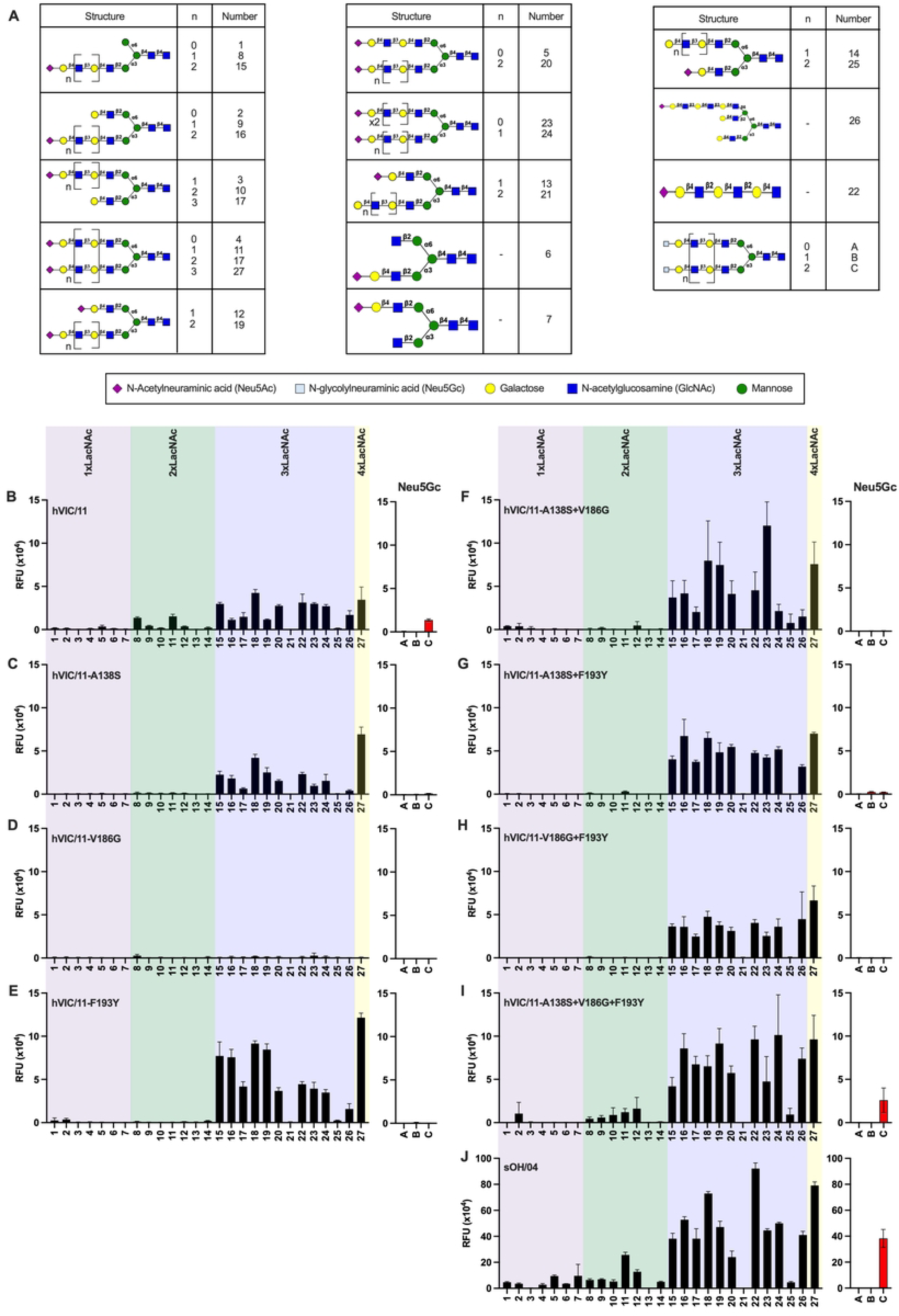
Swine-adaptative mutation limit the pool of α2,6 glycans supporting FLUAV binding. **a.** Structure of glycans assessed on the array. Mutant viruses (**b-i**) binding preference for different α2,6 glycans was assessed by glycan array. Viruses were normalized to 8 HAU and binding was performed for 1 hour at room temperature. Bound virus was detected using an anti-H3 stalk specific antibody. sOH/04 was used as a swine-adapted virus control (**j**). Binding to α2,6 Neu5Gc is shown in a separate panel (structures A, B, and C). Data is shown as relative fluorescent units (RFU) ± SD.

### Adaptative HA mutations enhance binding to extended α2,6-linked sialoglycans and increase α2,6-SA avidity

The A138S, V186G, and F193Y mutations are near the HA RBS but do not directly interact with sialic acid. Given their proximity to the RBS pocket, we tested virus binding to relevant complex *N*-glycans bearing terminal α2,6-, α2,3-linked Neu5Ac or Neu5Gc **(Fig 4A, S4 Fig)**. No binding was observed to α2,3-linked Neu5Ac or α2,3 Neu5Gc structures, regardless of the number of LacNAc repeats **(S4 Fig)**, highlighting a strict preference for α2,6-linked sialoglycans. All mutant viruses retained binding to α2,6 Neu5Ac structures, showing no preference for mono-, bi-, or tri-antennary structures **(Fig 4B-I)**. Parental hVIC/11 HA required at least 2 LacNAc repeats for binding and recognized α2,6-linked Neu5Gc only when 3 repeats were present. **(Fig 4B)**. Introduction of A138S prevented binding to glycans with less that 3 LacNAc repeats and completely abolished Neu5Gc recognition **(Fig 4C)**. Contrary, V186G displayed minimal binding across all structures tested **(Fig 4D),** consistent with its inability to agglutinate RBCs, whereas F193Y showed a similar pattern as A138S **(Fig 4E)**. Double mutants mirrored the binding pattern of A138S and F193Y (**Fig 4F-H**) Strikingly, the triple mutant (A138S+V186G+F193Y) regained broad α2,6-linked Neu5Ac binding, including glycans with a single LacNAc moiety, and restored Neu5Gc binding (**Fig 54I)**. Similarly, sOH/04 bound extensively to α2,6-linked Neu5Ac structures regardless of the amount of LacNAc repeats, including α2,6-linked Neu5Gc **(Fig 4J)**.

To complement these findings, receptor avidity was evaluated by biolayer interferometry (BLI). Streptavidin coated probes were loaded with either α2,3 sialyl LacNAc-PAA-biotin (α2,3, 3’SNL) or α2,6 sialyl LacNAc-PAA-biotin (α2,6, 6’SNL), viruses were normalized to 50 pM, and virus binding was evaluated in presence of oseltamivir carboxylate (OC, 10 μM) to prevent sialic acid hydrolysis. Consistent with the glycan array data, hVIC/11 and all the variants showed minimal binding to 3’SNL, whereas their binding for 6’SLN varied (**Fig 5A-H)**. Similarly, sOH/04 bound poorly to 3’SLN and exhibited a preference for 6’SLN **(Fig 5I)**. All mutants displayed higher 6’SNL responses than hVIC/11, with A138S and F193Y showing a 3-fold increase in binding and no statistical differences compared to sOH/04 **(Fig 5J)**. Combination of two or more mutations further enhanced binding for 6’SLN, confirming that these changes individually and collectively increase binding for α2,6-linked SA. Together, these findings show that HA mutations acquired during adaptation to swine under immune pressure narrow the spectrum of glycans recognized by the viruses, even though they do not directly interact with sialic acid. Additionally, these HA changes seem to enhance avidity for α2,6-SA.

**Fig 5.**
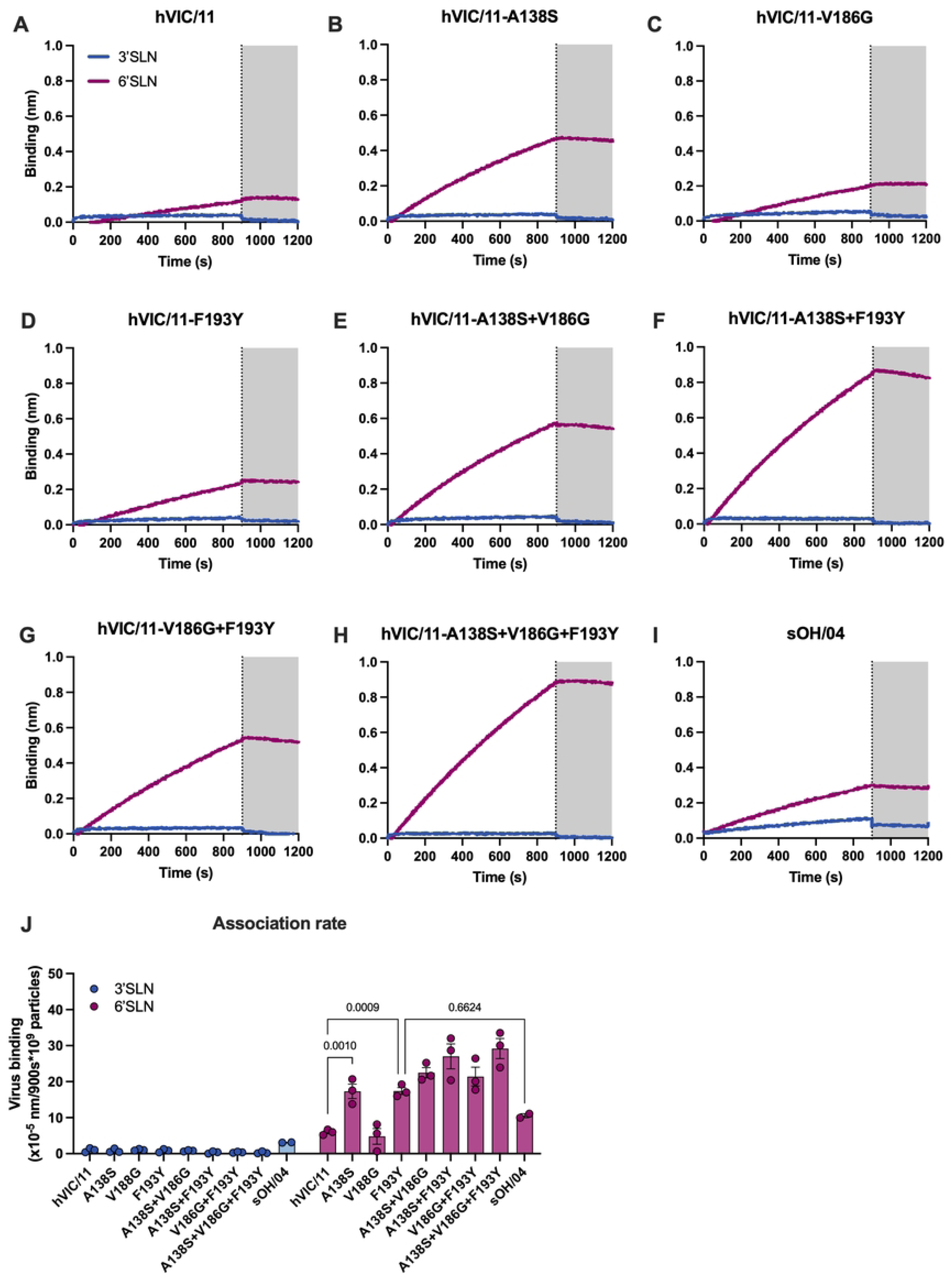
HA mutations enhance a2,6 SA binding. **a.** Binding strength for 3’SLN (blue) and 6’SLN (purple) was determined by BLI. Sensors were loaded with each polymer for 600 s. Excess polymer was washed away and binding of 50 nM hVIC/11 (**b**), hVIC/11 HA variants (**c-i**), and the control sOH/04 (**j**) was performed for 900 s in presence of oseltamivir carboxylate. Virus elution in presence of oseltamivir was performed for 300 s. Shaded are represent the dissociation step. Association rate (**k**) was calculated by determining the slope of each curve and normalized to the 10^9^ viral particles. Data is presented as the mean of least two independent experiments ± SD.

### The NA D113A mutation is not required to maintain the HA/NA balance when F193Y is present in the HA

We have previously shown that the NA D113A mutation compensates for the HA A138S substitution by disrupting calcium coordination in the four-fold symmetry axis, which decreases NA activity and restores the HA/NA balance. [32]. This is consistent with our data from non-vaccinated pigs in which this was the only mutation observed. In vaccinated animals, the D113A mutation was detected as a major variant only in C1 pigs. To determine whether its loss was due to a disturbed HA/NA balance with other HA mutants, we performed BLI experiments by normalizing viruses to 50 pM.

The sOH/04 control virus required 804 s to elute 50% of bound virus **(Fig 6A)**, whereas the hVIC/11 eluted more rapidly, regardless of D113A presence (190 s WT NA; 125 s D113A NA) **(Fig 6B)**. The hVIC/11-A138S showed prolonged elution times (478 s) with WT NA compared to hVIC/11, which further increased to 736 s upon introduction of D113A **(Fig 6C)**, resembling sOH/04. Introduction of D113A to hVIC/11-V186G resembled the effect observed for hVIC/11 **(Fig 6D)**. The F193Y mutant displayed elution kinetics similar to sOH/04 even in absence of D113A (688 s), and the D113A mutation had little effect **(Fig 6E)**. When multiple HA mutations were combined, the impact of D113A became minimal **(Fig 6F-I).** The triple HA mutant (A138S+V186G+F193Y) showed the slowest self-elution (>1,800 s), which seems to further increase with D113A **(Fig 6J),** suggesting that this combination particularly perturb the HA/NA balance. Overall, these results show that while D113A generally reduces virus self-elution, its compensatory role diminishes as HA binding to α2,6-SA increases. Additionally, this mutation is unnecessary in presence of F193Y, which intrinsically maintains the HA/NA balance to similar levels as sOH/04.

**Fig 6.**
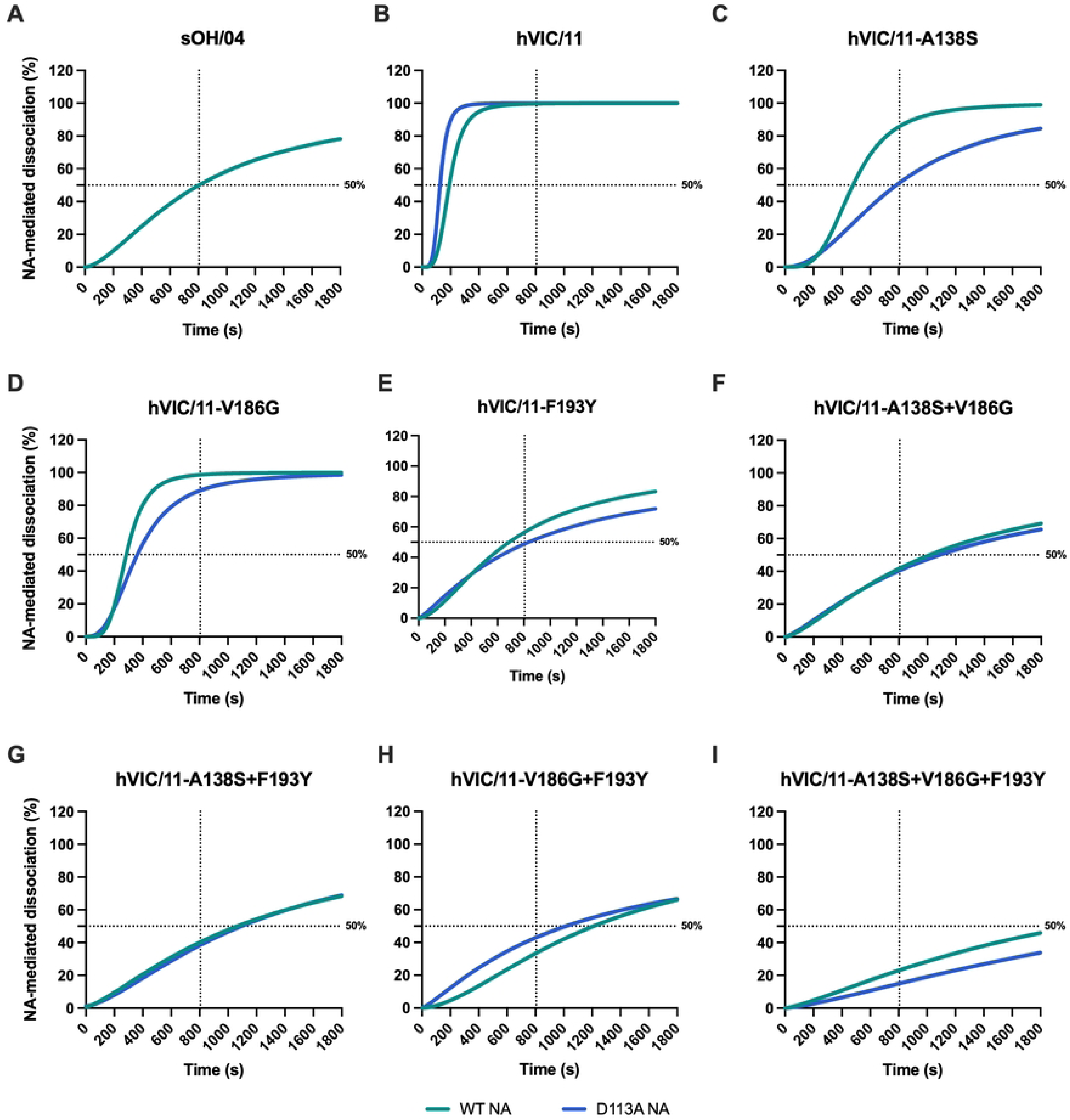
F193Y does not require NA changes to maintain the HA/NA balance. HA/NA balance of viruses carrying an hVIC/11 WT NA (green) or a D113A NA (blue) was evaluated by BLI. 50 nM of hVIC/11 (a), hVIC/11 variants (**b-h**), and sOH/04 (**i**) were bound to sensors previously loaded with 6’SLN for 900 s in presence of oseltamivir. Excess virus was washed away for 10 s in PBS containing oseltamivir, followed by oseltamivir removal by washing three times in PBS for 10 s each wash. NA-mediated virus dissociation was measured for 1,800 s. Elution time in the presence of NA activity was determined by fitting the data to a nonlinear four parameter variable slope equation and was defined as the time in which 50% of the virus was released from the sensor. Horizontal dotted line represents 50% elution while the vertical dotted line marks the time it took sOH/04 to elute 50% of virus (804 s). Data is presented as the mean of at least two independent experiments ± SD.

## DISCUSSION

Influenza A viruses contribute to the porcine respiratory disease complex and cause significant economic losses to the swine industry [33]. Vaccination remains the most effective control strategy, yet all currently licensed vaccines for use in swine in the United States are inactivated or vectored products [34]. These vaccines elicit strong antibody response against HA and NA that limits but might not prevent infection [35]. Vaccine-induced immunity exerts selective pressure that promotes antigenic drift and the selection of escape variants [11, 15, 16]. Such mutations not only reduce antibody recognition but may also drive evolution in a new host. However, despite the frequent spillover of human FLUAV into vaccinated swine herds, the role of prior immunity to endemic swine FLUAV on the evolution of human-origin viruses within pigs is unknown.

Mutations in the HA RBS are critical for host adaptation, particularly for avian viruses that must shift receptor specificity from α2,3- to α2,6-linked SA to infect pigs [36–38]. However, human seasonal H3N2 viruses, which prefer α2,6-SA, still replicate poorly in pigs despite similar α2,6-linked sialoglycans distribution in both species [39] [29]. Using our established *in vivo* transmission model [28, 29], we examined how pre-existing vaccine immunity shapes evolution of a human-like H3N2 virus (hVIC/11) in pigs. Vaccinated animals mounted a strong antibody response that limited replication but did not block transmission of hVIC/11-A138S, indicating that vaccine-induced immunity may not prevent human-to-swine spillovers. Deep sequencing revealed no HA changes in non-vaccinated animals, while in vaccinated pigs S138 rapidly reverted to A138, accompanied by the emergence of two additional RBS mutations, V186G and F193Y. Notably, only F193Y became fixed over 4 transmission events. Located within the antigenic site B, mutations at position 193 are known to promote antigenic drift in human and swine H3s [40–42]. F193Y has also been shown to enhance binding and replication of a human H3N2 virus in swine tracheal cells to levels comparable to A138S [29]. In our study, hVIC/11-F193Y exhibited the lowest HI titers among all variants and likely replicated in the lungs, as evidenced by detectable titers in BALF at the end of the experiment. These findings suggest that F193Y was selected as a variant combining immune evasion and efficient replication in swine. Similarly, residue 186 has also been associated with antigenic drift in human H3N2 viruses [43, 44], further reinforcing the idea that these variants emerged as a consequence of immune pressure.

Serologic data indicated that vaccine-induced immunity primarily targeted the HA, as HI titers were substantially higher than NI titers. Our NI assay uses the small substrate MUNANA, which can still access the catalytic site if antibodies do not directly block it. With this in mind, these results suggest that vaccine-induced antibodies did not target the NA substrate-binding site. The D113A mutation increased NI titers in most of the viruses we tested, regardless of the HA mutations. Residue 113 is located in the four-fold symmetry axis of N2s and modulates tetramer stability by preventing calcium coordination in the low-affinity calcium-binding pocket, a mechanism we have previously shown restores the HA/NA balance [45]. Perturbations at this site (e.g. antibody binding) could destabilize the tetramer, indirectly increasing NI titers even if antibodies are not targeting the catalytic site. Further evidence beyond the scope of this study is needed to support this hypothesis.

Receptor specificity strongly affects FLUAV interspecies transmission, yet the glycan preferences of human versus swine viruses remain poorly understood [19, 46, 47]. Mutations at positions 138, 186, and 193, which do not interact with sialic acid, greatly altered receptor binding. Despite pigs having higher α2,3-SA content in the lower respiratory tract (compared to humans), none of the variants bound α2,3 glycans, indicating that adaptation at the human-swine interface is not solely determined by sialic acid linkage. Notably, hVIC/11 bound Neu5Gc structures, but swine-adaptative mutations (A138S, V186G, and F193Y) abolished this recognition, suggesting that Neu5Gc is unlikely to play a major role during early adaptation of human H3N2 viruses to pigs. However, due to the limited number of Neu5Gc structures tested here, more studies will be needed to confirm this.

Glycan topology has been proposed as a determinant of FLUAV adaptation [48], with human H3N2 viruses evolving to recognize branched, extended structures [49]. We found that hVIC/11 bound glycans with at least 2 LacNAc repeats, whereas viruses carrying swine-adaptative mutations required at least 3, suggesting that adaptation enhanced binding to extended structures. Residues outside the canonical sialic acid binding pocket have been reported to participate in glycan binding [50]. Contemporary human H3N2s engage antigenically relevant residues in binding by forming hydrogen bonds along the glycan chain [50]. Our data supports this mechanism, as F193Y, an escape mutant, also influenced glycan binding. This is notable given that poly-LacNAc repeating glycans are particularly rare in swine lungs [51]. Although these mutations narrowed the pool of receptors recognized, they simultaneously enhanced avidity. This suggests a trade-off between receptor breadth and avidity, favoring multivalent interaction that could engage multiple receptor-binding sites within a single HA trimer as previously proposed [49]. A similar phenomenon has been observed at the human-avian interface, in which glycans that do not support binding on their own can compensate for the low abundance of preferred receptors by promoting hetero-multivalent interactions [52].

Finally, assessment of the HA/NA balance showed that the parental hVIC/11 eluted rapidly regardless of its NA background, consistent with weak receptor binding. In contrast, hVIC/11-A138S exhibited prolonged elution that was further extended by D113A, bringing balance to levels observed for sOH/04. The F193Y mutant displayed elution kinetics comparable to sOH/04 even without D113A, and introduction of D113A had little effect. This drastically contrast with V186G, which eluted very quickly, most likely due to low receptor binding combined with high NA activity. These findings explain why V186G was rapidly outcompeted and indicate that F193Y intrinsically maintains an optimal HA/NA balance and low neutralization by antibodies.

Overall, our findings demonstrate that vaccine-induced immunity in pigs can shape the evolution of human-origin H3N2 viruses. This process seems to select mutants with restricted receptor binding but enhanced α2,6-SA avidity while maintaining an optimal HA/NA balance. The fixation of F193Y highlights how immune escape and host adaptation can converge to promote viral fitness in a new host. These results indicate that vaccination may inadvertently accelerate viral adaptation following a spillover by favoring mutation of antigenically relevant residues that can alter glycan selectivity and enhance binding to host-specific structures.

## MATERIALS AND METHODS

### Ethics statement

All animal experiments presented in this study were reviewed and approved by the Institutional Care and Use Committee (IACUC) at the University of Georgia (Protocol #A2019 03-031-Y3-A9). Animals were housed under biosafety level 2 containment and cared for following the guidelines in the Guide for Care and Use of Agricultural Animals in Research and Teaching (American Dairy Science Association®, American Society of Animal Science, and Poultry Science Association, 2020). Water and feed were provided *ad libitum*. Animals were euthanized at the end of the study according to the guidelines provided by the American Veterinary Medical Association (AVMA).

### Cells and viruses

Madin-Darby canine kidney (MDCK), and Human Embryonic Kidney 293 (HEK-293) cells were maintained in Dulbecco’s Modified Eagles Medium (DMEM, Sigma-Aldrich, St Louis, MO) supplemented with 2mM L-glutamine (Sigma-Aldrich, St Louis, MO), 10% fetal bovine serum (FBS, Sigma-Aldrich, St Louis, MO), and 1% antibiotic/antimycotic (Sigma-Aldrich, St Louis, MO). HEK-293 and MDCK cells expressing A/WSN PB1 (HEK-293-PB1 and MDCK-PB1) were cultured as previously described [53]. HEK-293-PB1 were maintained in the media described above while MDCK-PB1 cells were grown in DMEM supplemented with 2mM L-glutamine (Sigma-Aldrich, St Louis, MO), 10% fetal bovine serum (FBS, Sigma-Aldrich, St Louis, MO), 1% antibiotic/antimycotic (Sigma-Aldrich, St Louis, MO), 0.25 μg/mL Puromycin (ThermoFisher Scientific, Waltham, MA), and 1 mg/mL Geneticin (ThermoFisher Scientific, Waltham, MA). Cells were incubated at 37°C in a humidified incubator with 5% CO_2_.

Human Airway epithelial cells (BCi.NS1.1) were kindly provided by Dr. Ronald Crystal (Weill Cornell Medicine, NY, USA) [54]. They were cultured in PneumaCult-Ex Plus Basal Media (STEMCELL Technologies, Vancouver, Canada) supplemented with PneumaCult-Ex Plus Supplement (STEMCELL Technologies, Vancouver, Canada), 0.1% hydrocortisone (STEMCELL Technologies, Vancouver, Canada), 1% antibiotic/antimycotic (Sigma-Aldrich, St Louis, MO), and 0.5% gentamycin (Sigma-Aldrich, St Louis, MO). For differentiation, cells were plated in type IV collagen-coated 12 mm transwell plates (0.4 μm pore size, Corning Inc., NY, USA) at a density of 3x10^5^ cells/well. After reaching 100% confluency, media for the apical compartment was removed and cells were incubated with PneumaCult ALI Base Media (STEMCELL Technologies, Vancouver, Canada), supplemented with PneumaCult ALI Supplement (STEMCELL Technologies, Vancouver, Canada), 1% PneumaCult ALI Maintenance Supplement (STEMCELL Technologies, Vancouver, Canada), 1% antibiotic/antimycotic (Sigma-Aldrich, St Louis, MO), 0.5% gentamycin (Sigma-Aldrich, St Louis, MO), 0.2% heparin (STEMCELL Technologies, Vancouver, Canada), and 0.5% hydrocortisone (STEMCELL Technologies, Vancouver, Canada). After 5 days of incubation at 37°C in a humidified incubator with 8% CO_2,_ cells were moved to an incubator at 37°C with 5% CO_2_ for 16 more days.

Viruses used in this study **(Table 1)** were grown in MDCK cells using Opti-MEM (ThermoFisher Scientific, Waltham, MA) containing 1 μg/ml of tosylsulfonyl phenylalanyl chloromethyl ketone (TPCK)-treated trypsin (Worthington Biochemicals, Lakewood, NJ). For biolayer interferometry experiments, replication-incompetent versions of the viruses were used. These viruses lacked the PB1 open reading (ΔPB1) and therefore could only infect and replicate in MDCK- and HEK-293T-PB1 cells. ΔPB1 viruses were grown in MDCK-PB1 cells using Opti-MEM (ThermoFisher Scientific, Waltham, MA) containing 1 μg/ml of TPCK-treated trypsin (Worthington Biochemicals, Lakewood, NJ), 0.25 μg/mL Puromycin (ThermoFisher Scientific, Waltham, MA), and 1 mg/mL Geneticin (ThermoFisher Scientific, Waltham, MA) as previously described [53]. Infections were incubated at 37°C for 3 days and viral titers were determined by TCID_50_ using the Reed and Muench method [56].

**Table 1.** Viruses used in this study*.

| Abbreviation | Strain | Surface segments | Backbone |
| --- | --- | --- | --- |
| hVIC/11* | rgA/Victoria/361/2011 | hVIC/11 HA and NA | TRIG |
| hVIC/11-A138S* | rgA/Victoria/361/2011-A138 | hVIC/11 HA A138S and NA | TRIG |
| hVIC/11-V186G* | rgA/Victoria/361/2011-V186G | hVIC/11 HA V186G and NA | TRIG |
| hVIC/11-F193Y* | rgA/Victoria/361/2011-F193Y | hVIC/11 HA F193Y and NA | TRIG |
| hVIC/11-A138S+V186G* | rgA/Victoria/361/2011-A138S+V186G | hVIC/11 HA A138S+V186G and NA | TRIG |
| hVIC/11-A138S+F193Y* | rgA/Victoria/361/2011-A138S+F193Y | hVIC/11 HA A138S+F193Y and NA | TRIG |
| hVIC/11-A138S+V186G+F193Y* | rgA/Victoria/361/2011-A138S+V186G+F193Y | hVIC/11 HA A138S+V186G+F193Y and NA | TRIG |
| hVIC/11-D113A* | rgA/Victoria/361/2011-D113A | hVIC/11 HA and NA D113A | TRIG |
| hVIC/11-A138S/D113A* | rgA/Victoria/361/2011-A138/D113A | hVIC/11 HA A138S and NA D113A | TRIG |
| hVIC/11-V186G* | rgA/Victoria/361/2011-V186G | hVIC/11 HA V186G and NA D113A | TRIG |
| hVIC/11-F193Y* | rgA/Victoria/361/2011-F193Y/D113A | hVIC/11 HA F193Y and NA D113A | TRIG |
| hVIC/11-A138S+V186G* | rgA/Victoria/361/2011-A138S+V186G/D113A | hVIC/11 HA A138S+V186G and NA D113A | TRIG |
| hVIC/11-A138S+F193Y/D113A* | rgA/Victoria/361/2011-A138S+F193Y/D113A | hVIC/11 HA A138S+F193Y and NA D113A | TRIG |
| hVIC/11-A138S+V186G+F193Y/D113A* | rgA/Victoria/361/2011-A138S+V186G+F193Y/D113A | hVIC/11 HA A138S+V186G+F193Y and NA D113A | TRIG |
| sOH/04* | rgA/turkey/Ohio/313053/2004 | sOH/04 HA and NA | TRIG |
| NY/11 | rgA/swine/New York/A01104005/2011 | NY/11 HA and NA | Wild type |

\*rg backbone derived from sOH/04 triple reassortant (TRIG)[55]

### Animal studies

Three-week-old cross-bred pigs were obtained from Midwest Research Swine Inc (Glencoe, MN, USA). After a 7-day acclimatation period, animals were confirmed to be seronegative for anti-FLUAV antibodies by competitive ELISA (IDEXX, Westbrook, ME) according to the manufacturer’s instructions. Pigs were randomly assigned into four experimental group. Animals in two groups (n=15/group) were immunized intramuscularly using the Zoetis FluSure XP vaccine (Zoetis, Parsippany, NJ) according to the manufacturer’s instructions. Animals in two groups (n=15/group) were mock vaccinated with PBS. Fourteen days post-vaccination (dpv), primed pigs were boosted using the same vaccine. Fourteen days post-boost (dpb), blood was collected to confirm seroconversion induced by the vaccine. Three pigs/group (seeders) were sedated with ketamine (6 mg/kg, Zoetis, Parsippany, NJ), xylazine (3 mg/kg, Cronus Pharma, Brunswick, NJ), and Telazol (6 mg/kg, Dechra Pharmaceutical, Fort Worth, TX), and challenged intratracheally (2ml) and intranasally (1ml) with 3x10^6^ TCID_50_/pig of hVIC/11-A138S [28]. At 2 dpi, same-vaccination-status pigs (n=3, Contact 1 [C1]) were placed in contact with the directly-inoculated animals (seeders). At 5 dpi, seeders were sedated using a mixture of ketamine (3 mg/kg), xylazine (1.5 mg/kg), and Telazol (3 mg/kg) and then humanely euthanized by an intravenous pentobarbital overdose (Euthasol, 390 mg/10, Virbac, Westlake, TX). C1 pigs were moved to a clean cage and 3 new, same-vaccination-status pigs (C2) were put in contact with C1 pigs. At 5 dpi, C1 pigs were sedated and humanely euthanized as described above. This cycle was repeated for a total of four contacts. Pigs were checked daily for clinical signs and nasal swabs were collected at 0, 1, 2, 3, and 5 days post-infection (dpi) or 0, 1, 3, 5, and 6 days post-contact (DPC) in 2 mL of brain heart infusion broth (Sigma-Aldrich, St Louis, MO). Anti-FLUAV antibody levels were measured for every contact the day they were place in contact with infected animals. During necropsies, lungs were rinsed with 50 mL of DMEM to collect the bronchoalveolar lavage fluid (BALF).

### Enzyme-linked immune absorbent assay (ELISA)

To quantify antibody levels induced by the Zoetis FluSure XP vaccine, an in-house ELISA was performed: the vaccine contains two H3 components (1990.4.a and 1990.4.b HA clades) along with two H1 components. Briefly, high-binding ELISA plates (ThermoFisher Scientific, Waltham, MA) were coated with 128 HAU of rg-A/swine/NY/A01104005/2011 (H3N2, 1990.4.a) diluted in ELISA coating buffer (ThermoFisher Scientific, Waltham, MA). The virus was previously purified by ultracentrifugation using a 20% sucrose cushion at 30,000 rpm. After a 2-hour adsorption, plates were washed twice with washing buffer (ThermoFisher Scientific, Waltham, MA) and 100 μL of 1:10 diluted serum sample was added and incubated for 1 hour at room temperature. After incubation, plates were washed three times with washing buffer and an HRP-conjugated anti-swine IgG secondary antibody was added (ThermoFisher Scientific, Waltham, MA) in a 1:1000 dilution. The secondary antibody was incubated for one hour, and excess antibody was washed away three times with washing buffer. Finally, 100 μL of 3,3’,5,5’-tetramethylbenzidine (TMB, ThermoFisher Scientific, Waltham, MA) was added per well and plates were incubated for 10 minutes in the dark. Reaction was stopped by adding 50 μL of stop solution (ThermoFisher Scientific, Waltham, MA) and absorbance was measured at 450 nm using a Synergy HTX Multi-Mode Microplate Reader (Agilent BioTek, Santa Clara, CA). Samples with an absorbance 3s higher than the mean of mock-vaccinated pigs were considered positive.

### Nasal swabs and BALF virus titration

vRNA from nasal swabs and BALF samples was extracted using the MagMax-96 AI/ND viral RNA isolation kit (ThermoFisher Scientific, Waltham, MA) according to the manufacturer’s instructions. Purified RNA was used to titrate FLUAV using qRT-PCR using the Quantabio qScript XLT One-Step RT-qPCR ToughMix kit (Quantabio, Beverly MA) targeting the M segment as previously described [32].

### Next-generation sequencing (NGS)

Purified RNA was used as template to amplify the HA and NA segments with target-specific primers and using the SuperScript III One-Step PCR System (ThermoFisher Scientific, Waltham, MA) according to the manufacturer’s instructions. Amplification was confirmed by running the PCR products in 1% agarose gels. Replicate HA and NA reactions were then combined and purified using 0.4X Agencourt AMPure XP Magnetic Beads (Beckman Coulter, Brea, CA, USA). DNA concentration was determined using the Qubit dsDNA HS assay kit (ThermoFisher Scientific, Waltham, MA) on a Qubit 3.0 fluorometer (ThermoFisher Scientific, Waltham, MA) and normalized to 0.2 ng/μL. PCR products were fragmented and indexed using the Nextera XT DNA library preparation kit (Illumina, San Diego, CA, USA). Tagmentation was confirmed by running random samples on an Agilent Bioanalyzer 2100 DNA-HS assay (Agilent, Santa Clara, CA, USA). Finally, samples were normalized to 0.2 ng/μL, pooled, denatured, normalized to 10 pM, and sequenced using the MiSeq Reagent Kit V2, 300 cycles (Illumina, San Diego, CA, USA).

### Variant analysis

Variant analysis was performed as previously described [32, 57]. Briefly, Adapters were removed using Cutadapt 3.4 and mapped to a reference sequence using *mem* from BWA version 0.7.17 [58]. Non consensus variants were identified using LoFreq [59] following the best practices stablished in the Genome Analysis Toolkit [60]. Only variants with a frequency of 1% and a minimum coverage of 100 were used. Synonymous and nonsynonymous variants were identified using SNPdat version v1.0.5 [61]. Variants with a frequency above 50% were considered major variants.

### Site-directed mutagenesis and virus rescue

Mutations were introduced in the HA and NA genes of rg-A/Victoria/361/2011 using the Phusion site-directed mutagenesis kit (ThermoFisher Scientific, Waltham, MA) according to the manufacturer’s instructions and plasmid sequences were confirmed by whole plasmid sequencing. Recombinant viruses were rescued using an 8-plasmid reverse genetic system using a coculture of HEK-293 and MDCK cells as previously described [62].

To generate replication-incompetent viruses, rescues were performed as described above using a coculture of HEK-293-PB1 and MDCK-PB1 cells without antibiotics. The pDP plasmid containing the TRIG PB1 gene was replaced with a pHW_PB1packWSN_TdKatushka NLS plasmid which contained the packaging signals of the PB1 segment and the ORF of the TdKatushka fluorescent protein instead of the PB1 coding sequence [63].

### *In vitro* growth kinetics

MDCK cells were seeded in 6-well plates at a density of 3x10^5^ cells/cm^2^ in Opti-MEM and incubated at 37°C in a humidified incubator with 5% CO_2_ until an 80% confluency before use. Cells were infected at a multiplicity of infection (MOI) of 0.01 for 1 hour at 37°C. After incubation, cells were washed three times with PBS and supplemented with Opti-MEM containing 1 μg/ml TPCK-treated trypsin. Supernatants were collected at 0, 12, 24, 48, and 72 hpi timepoints.

Differentiated HAE cells were infected at an MOI of 0.01 by adding 200 μL of inoculum in the apical compartment. After 1 hour, cells were washed 5 times with PBS and transwells were transferred to a new plate. Cells were maintained in ALI media and no exogenous TPCK-treated trypsin was added. Supernatants were collected at 0, 12, 24, 48, and 72 hpi by adding 200 mL of PBS onto the cells and incubating them at 37°C for 10 minutes. Viral RNA was extracted, and titers determined by qRT-PCR as described above and data was fit to a TCID_50_ equivalent standard curve of an exact virus match.

### Virus neutralization assay

Because hVIC/11-V186G does not agglutinate chicken nor turkey red blood cells (RBC), neutralization titers were determined using a homologous fluorescent virus to rely on mCherry expression in infected cells, similar to luciferase expression as previously described [64]. Briefly, viruses listed in **Table 1** were rescued as mentioned above in the isogenic TRIG backbone but carrying the mCherry coding sequence downstream of the NS1 gene. Three representative sera samples were treated with receptor-destroying enzyme (RDE, Denka Seiken, VWR) at 37°C for 18 hours and then RDE was heat-inactivated for 30 minutes at 56°C. Sera was diluted 1:10 and further mixed in a 1:1 ratio with 100 TCID_50_ of each virus for 1 hour at 37°C. Then the mixture was added onto confluent MDCK cells in 96 well-plates. Infections were incubated for 1 hour at 37°C and then cells were washed three times with PBS and overlayed with Opti-MEM supplemented with 1 μg/ml TPCK-treated trypsin. At 48 hours post infection, cell culture media was removed, and cells were lysed using the Pierce IP lysis buffer (ThermoFisher Scientific, Waltham, MA). Supernatant was collected and mCherry fluorescence was measured at excitation and emission wavelengths of 587 nm and 610 nm, respectively, using a Synergy HTX Multi-Mode Microplate Reader (Agilent BioTek, Santa Clara, CA).

### Hemagglutinin inhibition assays

Three representative sera samples were treated with RDE as described above. Samples were 2-fold serially diluted in PBS and then 25 μL of each sera samples were mixed with 25 μL of virus containing 4 hemagglutination units (HAU) in V-bottom 96-well plates. Sera/virus mixtures were incubated for 30 minutes at room temperature and after incubation 50 μL of 0.5% turkey red blood cells were added. Following a 40-minute incubation at room temperature, HI titers were determined and titers below 10 HI were arbitrarily assigned a value of 10.

### Neuraminidase Inhibition assays

Three representative sera samples were subjected to RDE treatment as described above. Anti-sera was then 2-fold serially diluted in reaction buffer (32.5 mM 2-(N-morpholino)ethanesulfonic acid (MES), 2 mM CaCl_2_, pH 6.5). At the same time, viruses were normalized based on NA activity as previously described [65]. Subsequently, 50 μL of anti-sera were mixed at a 1:1 ratio with virus dilution and incubated for 30 minutes at room temperature.

Neuraminidase activity was evaluated as previously described [32, 65]. Briefly, after the 30 minutes incubation, 50 μL of MUNANA were added to each sample to a final concentration of 100 μM MUNANA. 4-methylumbelliferone (4-MU) production was monitored by measuring fluorescence every 60 seconds at excitation and emission wavelengths of 360 nm and 460 nm respectively, using a Synergy HTX Multi-Mode Microplate Reader (Agilent BioTek, Santa Clara, CA).

### Glycan array

Array experiments were performed as previously described [66]. Briefly, viruses were diluted to 8 HAU in TSM buffer (20 mM Tris-HCl, 150 mM NaCl, 2 mM CaCl_2_, and 2 mM MgCl_2_, pH 7.4). 100 μL of each sample was applied onto the slide and viruses were incubated for 1 hour at room temperature in presence of 10 μM Oseltamivir Carboxilate (Sigma-Aldrich, St Louis, MO). After incubation, slides were washed three times with TSM-wash (TSM-buffer with 0.05% Tween-20) and one time with water. Slides were then incubated with 5 μg/mL of an anti-HA antibody (stem specific antibody, CR8020, Absolute Antibody, Shirley, MA) for one hour. Samples were washed three times as described above and then incubated with 5 μg/mL of AlexaFluor 647-labeled goat anti-human IgG secondary antibody (Jackson ImmunoResearch Cat #109-605-008). After a 1-hour incubation, slides were washed three times with TSM-wash, one time with water, and air dried. The slides were scanned using a GenePix 4000B microarray scanner (Molecular Devices) at the appropriate excitation wavelength with a resolution of 5 μm. Optimum gains and PMT values were employed for the scanning ensuring that all signals were within the linear range of the scanner’s detector and there was no saturation of signals. The images were analyzed using GenePix Pro 7 software (version 7.2.29.2, Molecular Devices). The data were analyzed with a home written Excel macro. The highest and the lowest value of the total fluorescence intensity of the replicate spots were removed, and the remaining values were used to provide the mean value and standard deviation. The fluorescence values were plotted using Prism 10 software (GraphPad Software, Inc.), bars represent the mean ± SD for each treatment.

### Nanoparticle tracking analysis (NTA)

Viral particle concentration of replication-incompetent virus preparations was quantified using a NanoSight NS300 nanoparticle tracker analyzer (Malvern, UK) as previously described [67]. Briefly, samples were diluted in ultrapure water to 10^8^-10^9^ particles/mL to fit the linear range of the instrument. Particle concentration was determined by recording 60 second sample videos, five independent times. Data was then analyzed using the Nanoparticle Tracker analysis 3.0 Software (Malvern, UK).

### Biolayer Interferometry (BLI) experiments and HA/NA balance assessment

Replication-incompetent viruses were normalized to 50 pM based on the NTA results and BLI experiments were performed as previously described [67]. Briefly, streptavidin sensors were loaded to saturation for 600 s with 50 μg /mL of either Neu5Acα2-3Galβ1-4GlcNAcβ-PAA-biotin (α2,3, 3’SNL, Glycotech, Gaithersburg, MD) or Neu5Acα2-6Galβ1-4GlcNAcβ-PAA-biotin (α2,6, 6’SNL, Glycotech, Gaithersburg, MD) diluted in reaction buffer (PBS supplemented with 1% BSA, 0.1% Tween-20, and 2 mM CaCl_2_, pH 6.5). Excess receptor was washed away for 300 s. Virus association was performed for 900 s in presence of 10 μM oseltamivir (Sigma-Aldrich, St Louis, MO) followed by a 300 s dissociation step in presence of oseltamivir. Binding curves were plotted, and the association rate was determined as previously described [67]. Binding of our FLUAV viruses to 6’SLN resulted in virtually irreversible attachment (>10% dissociation). This prevented calculation of *k_off_* and therefore K_D_ could not be determined. To analyze the HA/NA balance, virus association was performed as described above followed by 3x10 s washes in reaction buffer without oseltamivir. Finally, NA-mediated dissociation was measured for 1,800 s in absence of oseltamivir. All measurements were performed using an Octet RED96e System (ForteBio, Dallas, TX). Data acquisition and evaluation was performed using the ForteBio Data Acquisition 11.1 and ForteBio Data Analysis 11.1 software, respectively. Response curves raw data was exported, and curves were flipped i.e., corrected to give positive response (by multiplying responses with -1). Curves depicting virus binding (thickness, nm) vs. time (sec) were plotted using the Prism 10 software (GraphPad Software, Inc.). The assay was performed at least two times for each virus. Finally, NA-mediated dissociation was determined as the relationship between the response observed at any given time compared to the maximum binding.

### Statistics and reproducibility

All statistical analysis presented here were performed using GraphPad Prism Version 10.4.1. P values were obtained by ordinary one-way ANOVA with Tukey’s multiple comparison test, and no data was excluded from the analysis. *In vitro* analyses were performed at least three times unless stated otherwise in the figure legend, and results were reproducible between experiments.

## ACKNOWLEDGMENT

We thank Adrian Creanga and Masaru Kanekiyo for kindly providing the plasmids necessary to construct ΔPB1 viruses.

## REFERENCES

1. Nelson, M.I., et al., Global transmission of influenza viruses from humans to swine. J Gen Virol, 2012. 93(Pt 10): p. 2195–2203.

2. Nelson, M.I., et al., Introductions and evolution of human-origin seasonal influenza a viruses in multinational Swine populations. J Virol, 2014. 88(17): p. 10110–9.

3. Nelson, M.I. and A.L. Vincent, Reverse zoonosis of influenza to swine: new perspectives on the human-animal interface. Trends Microbiol, 2015. 23(3): p. 142–53.

4. Rajao, D.S., A.L. Vincent, and D.R. Perez, Adaptation of Human Influenza Viruses to Swine. Front Vet Sci, 2018. 5: p. 347.

5. Rajao, D.S., et al., Changes in the Hemagglutinin and Internal Gene Segments Were Needed for Human Seasonal H3 Influenza A Virus to Efficiently Infect and Replicate in Swine. Pathogens, 2022. 11(9).

6. Nelson, M.I., et al., Evolution of novel reassortant A/H3N2 influenza viruses in North American swine and humans, 2009-2011. J Virol, 2012. 86(16): p. 8872–8.

7. Nelson, M.I., et al., Genomic reassortment of influenza A virus in North American swine, 1998-2011. J Gen Virol, 2012. 93(Pt 12): p. 2584–2589.

8. Kida, H., K.F. Shortridge, and R.G. Webster, Origin of the hemagglutinin gene of H3N2 influenza viruses from pigs in China. Virology, 1988. 162(1): p. 160–6.

9. Lewis, N.S., et al., Substitutions near the hemagglutinin receptor-binding site determine the antigenic evolution of influenza A H3N2 viruses in U.S. swine. J Virol, 2014. 88(9): p. 4752–63.

10. Vincent, A.L., et al., Influenza A virus vaccines for swine. Vet Microbiol, 2017. 206: p. 35–44.

11. Islam, S., et al., Influenza A haemagglutinin specific IgG responses in children and adults after seasonal trivalent live attenuated influenza vaccination. Vaccine, 2017. 35(42): p. 5666–5673.

12. Isakova-Sivak, I., et al., Influenza vaccine: progress in a vaccine that elicits a broad immune response. Expert Review of Vaccines, 2021. 20(9): p. 1097–1112.

13. Van Reeth, K., et al., Heterologous prime-boost vaccination with H3N2 influenza viruses of swine favors cross-clade antibody responses and protection. NPJ Vaccines, 2017. 2.

14. Murcia, P.R., et al., Evolution of an Eurasian avian-like influenza virus in naive and vaccinated pigs. PLoS Pathog, 2012. 8(5): p. e1002730.

15. Lopez-Valinas, A., et al., Evolution of Swine Influenza Virus H3N2 in Vaccinated and Nonvaccinated Pigs after Previous Natural H1N1 Infection. Viruses, 2022. 14(9).

16. Lopez-Valinas, A., et al., Identification and Characterization of Swine Influenza Virus H1N1 Variants Generated in Vaccinated and Nonvaccinated, Challenged Pigs. Viruses, 2021. 13(10).

17. Xiong, X.L., J.W. McCauley, and D.A. Steinhauer, Receptor Binding Properties of the Influenza Virus Hemagglutinin as a Determinant of Host Range. Influenza Pathogenesis and Control - Vol I, 2014. 385: p. 63–91.

18. Matrosovich, M., et al., Early alterations of the receptor-binding properties of H1, H2, and H3 avian influenza virus hemagglutinins after their introduction into mammals. J Virol, 2000. 74(18): p. 8502–12.

19. Stevens, J., et al., Receptor specificity of influenza A H3N2 viruses isolated in mammalian cells and embryonated chicken eggs. J Virol, 2010. 84(16): p. 8287–99.

20. Nicholls, J.M., et al., Sialic acid receptor detection in the human respiratory tract: evidence for widespread distribution of potential binding sites for human and avian influenza viruses. Respir Res, 2007. 8(1): p. 73.

21. Trebbien, R., L.E. Larsen, and B.M. Viuff, Distribution of sialic acid receptors and influenza A virus of avian and swine origin in experimentally infected pigs. Virol J, 2011. 8: p. 434.

22. Suzuki, T., et al., Swine influenza virus strains recognize sialylsugar chains containing the molecular species of sialic acid predominantly present in the swine tracheal epithelium. FEBS Lett, 1997. 404(2-3): p. 192–6.

23. Altman, M.O. and P. Gagneux, Absence of Neu5Gc and Presence of Anti-Neu5Gc Antibodies in Humans-An Evolutionary Perspective. Front Immunol, 2019. 10: p. 789.

24. Gambaryan, A.S., et al., Receptor-binding properties of swine influenza viruses isolated and propagated in MDCK cells. Virus Res, 2005. 114(1-2): p. 15–22.

25. Chien, Y.A., et al., Single Particle Analysis of H3N2 Influenza Entry Differentiates the Impact of the Sialic Acids (Neu5Ac and Neu5Gc) on Virus Binding and Membrane Fusion. J Virol, 2023. 97(3): p. e0146322.

26. Bradley, K.C., et al., Comparison of the receptor binding properties of contemporary swine isolates and early human pandemic H1N1 isolates (Novel 2009 H1N1). Virology, 2011. 413(2): p. 169–82.

27. Masuda, H., et al., Substitution of amino acid residue in influenza A virus hemagglutinin affects recognition of sialyl-oligosaccharides containing N-glycolylneuraminic acid. FEBS Lett, 1999. 464(1-2): p. 71–4.

28. Cardenas, M., et al., Amino acid 138 in the HA of a H3N2 subtype influenza A virus increases affinity for the lower respiratory tract and alveolar macrophages in pigs. PLoS Pathog, 2024. 20(2): p. e1012026.

29. Mo, J.S., et al., Transmission of Human Influenza A Virus in Pigs Selects for Adaptive Mutations on the HA Gene. J Virol, 2022. 96(22): p. e0148022.

30. Han, A.X., S.P.J. de Jong, and C.A. Russell, Co-evolution of immunity and seasonal influenza viruses. Nat Rev Microbiol, 2023. 21(12): p. 805–817.

31. Lee, P.S., et al., Receptor mimicry by antibody F045-092 facilitates universal binding to the H3 subtype of influenza virus. Nat Commun, 2014. 5: p. 3614.

32. Cardenas, M., et al., Modulation of human-to-swine influenza a virus adaptation by the neuraminidase low-affinity calcium-binding pocket. Commun Biol, 2024. 7(1): p. 1230.

33. Nakharuthai, C., et al., Occurrence of swine influenza virus infection in swine with porcine respiratory disease complex. Southeast Asian J Trop Med Public Health, 2008. 39(6): p. 1045–53.

34. Ma, W. and J.A. Richt, Swine influenza vaccines: current status and future perspectives. Anim Health Res Rev, 2010. 11(1): p. 81–96.

35. Lorbach, J.N., et al., Influenza Vaccination of Swine Reduces Public Health Risk at the Swine-Human Interface. mSphere, 2021. 6(3): p. e0117020.

36. Mancera Gracia, J.C., et al., Effect of serial pig passages on the adaptation of an avian H9N2 influenza virus to swine. PLoS One, 2017. 12(4): p. e0175267.

37. Yang, W., et al., Mutations during the Adaptation of H9N2 Avian Influenza Virus to the Respiratory Epithelium of Pigs Enhance Sialic Acid Binding Activity and Virulence in Mice. J Virol, 2017. 91(8).

38. Bateman, A.C., et al., Amino acid 226 in the hemagglutinin of H4N6 influenza virus determines binding affinity for alpha2,6-linked sialic acid and infectivity levels in primary swine and human respiratory epithelial cells. J Virol, 2008. 82(16): p. 8204–9.

39. Nelli, R.K., et al., Comparative distribution of human and avian type sialic acid influenza receptors in the pig. BMC Vet Res, 2010. 6: p. 4.

40. Abente, E.J., et al., The Molecular Determinants of Antibody Recognition and Antigenic Drift in the H3 Hemagglutinin of Swine Influenza A Virus. J Virol, 2016. 90(18): p. 8266–80.

41. Wang, X., et al., Amino Acids in Hemagglutinin Antigenic Site B Determine Antigenic and Receptor Binding Differences between A(H3N2)v and Ancestral Seasonal H3N2 Influenza Viruses. J Virol, 2017. 91(2).

42. Burke, D.F., Structural Consequences of Antigenic Variants of Human A/H3N2 Influenza Viruses. Viruses, 2023. 15(4).

43. Tewawong, N., et al., Assessing Antigenic Drift of Seasonal Influenza A(H3N2) and A(H1N1)pdm09 Viruses. PLoS One, 2015. 10(10): p. e0139958.

44. Lee, M.S. and J.S. Chen, Predicting antigenic variants of influenza A/H3N2 viruses. Emerg Infect Dis, 2004. 10(8): p. 1385–90.

45. Wang, H., et al., Structural restrictions for influenza neuraminidase activity promote adaptation and diversification. Nat Microbiol, 2019. 4(12): p. 2565–2577.

46. Zhao, C. and J. Pu, Influence of Host Sialic Acid Receptors Structure on the Host Specificity of Influenza Viruses. Viruses, 2022. 14(10).

47. Gambaryan, A., et al., Receptor specificity of influenza viruses from birds and mammals: new data on involvement of the inner fragments of the carbohydrate chain. Virology, 2005. 334(2): p. 276–83.

48. Chandrasekaran, A., et al., Glycan topology determines human adaptation of avian H5N1 virus hemagglutinin. Nat Biotechnol, 2008. 26(1): p. 107–13.

49. Peng, W., et al., Recent H3N2 Viruses Have Evolved Specificity for Extended, Branched Human-type Receptors, Conferring Potential for Increased Avidity. Cell Host Microbe, 2017. 21(1): p. 23–34.

50. Thompson, A.J., et al., Evolution of human H3N2 influenza virus receptor specificity has substantially expanded the receptor-binding domain site. Cell Host Microbe, 2024. 32(2): p. 261–275 e4.

51. Byrd-Leotis, L., et al., Shotgun glycomics of pig lung identifies natural endogenous receptors for influenza viruses. Proc Natl Acad Sci U S A, 2014. 111(22): p. E2241–50.

52. Liu, M., et al., Human-type sialic acid receptors contribute to avian influenza A virus binding and entry by hetero-multivalent interactions. Nat Commun, 2022. 13(1): p. 4054.

53. Creanga, A., et al., A comprehensive influenza reporter virus panel for high-throughput deep profiling of neutralizing antibodies. Nat Commun, 2021. 12(1): p. 1722.

54. Walters, M.S., et al., Generation of a human airway epithelium derived basal cell line with multipotent differentiation capacity. Respir Res, 2013. 14(1): p. 135.

55. Pena, L., et al., Modifications in the polymerase genes of a swine-like triple-reassortant influenza virus to generate live attenuated vaccines against 2009 pandemic H1N1 viruses. J Virol, 2011. 85(1): p. 456–69.

56. Reed, L.J. and H. Muench, A SIMPLE METHOD OF ESTIMATING FIFTY PER CENT ENDPOINTS12. American Journal of Epidemiology, 1938. 27(3): p. 493–497.

57. Ferreri, L.M., et al., Intra- and inter-host evolution of H9N2 influenza A virus in Japanese quail. Virus Evol, 2022. 8(1): p. veac001.

58. Li, H. and R. Durbin, Fast and accurate short read alignment with Burrows-Wheeler transform. Bioinformatics, 2009. 25(14): p. 1754–60.

59. Wilm, A., et al., LoFreq: a sequence-quality aware, ultra-sensitive variant caller for uncovering cell-population heterogeneity from high-throughput sequencing datasets. Nucleic Acids Res, 2012. 40(22): p. 11189–201.

60. McKenna, A., et al., The Genome Analysis Toolkit: a MapReduce framework for analyzing next-generation DNA sequencing data. Genome Res, 2010. 20(9): p. 1297–303.

61. Doran, A.G. and C.J. Creevey, Snpdat: easy and rapid annotation of results from de novo snp discovery projects for model and non-model organisms. BMC Bioinformatics, 2013. 14: p. 45.

62. Hoffmann, E., et al., Eight-plasmid system for rapid generation of influenza virus vaccines. Vaccine, 2002. 20(25-26): p. 3165–70.

63. Bloom, J.D., L.I. Gong, and D. Baltimore, Permissive secondary mutations enable the evolution of influenza oseltamivir resistance. Science, 2010. 328(5983): p. 1272-5.

64. Caceres, C.J., et al., Use of Reverse Genetics for the Generation of Recombinant Influenza Viruses Carrying Nanoluciferase. Methods Mol Biol, 2024. 2733: p. 47–74.

65. Marathe, B.M., et al., Determination of neuraminidase kinetic constants using whole influenza virus preparations and correction for spectroscopic interference by a fluorogenic substrate. PLoS One, 2013. 8(8): p. e71401.

66. Broszeit, F., et al., Glycan remodeled erythrocytes facilitate antigenic characterization of recent A/H3N2 influenza viruses. Nat Commun, 2021. 12(1): p. 5449.

67. de Vries, E., et al., Quantification of Receptor Association, Dissociation, and NA-Dependent Motility of Influenza A Particles by Biolayer Interferometry. Methods Mol Biol, 2022. 2556: p. 123–140.

